# Simultaneous TCR and IL-2 agonism selectively enhances epitope-specific CD8 T cell responses during chronic viral infection

**DOI:** 10.64898/2026.04.13.718150

**Authors:** Masao Hashimoto, Mohammad Affan Khan, Akil Akhtar, Javed N Agrewala, Gordon J. Freeman, Natasha Girgis, Yu Zhang, Simon Low, Steven N Quayle, Anish Suri, Rafi Ahmed

## Abstract

Interleukin-2 (IL-2) remains an attractive cytokine for enhancing antigen-specific CD8 T cell responses in cancer immunotherapy, but systemic toxicity hinders its broad clinical application. To address this, various IL-2-based therapeutics have been engineered with altered IL-2 receptor bias or targeted delivery to tumors, the tumor microenvironment, or immune cell populations. Ideally, IL-2 signals should be selectively delivered to antigen-specific CD8 T cells, boosting their responses and promoting effector differentiation while sparing non-targeted populations. Immuno-STAT^TM^ (Selective Targeting and Alteration of T cells) is a fusion protein platform comprising a bivalent peptide-MHC class I complex and an affinity-attenuated IL-2 mutein that co-stimulates TCR and IL-2 signaling in epitope-specific CD8 T cells. Here, we investigated whether a D^b^GP_33-41_-targeted Immuno-STAT enhances D^b^GP33-specific CD8 T cell responses in a mouse model of chronic lymphocytic choriomeningitis virus (LCMV) infection. Immuno-STAT treatment selectively expanded D^b^GP33-specific CD8 T cells with an effector-like phenotype. Non-targeted D^b^GP276-specific CD8 T cells showed little to no expansion in response to D^b^GP_33-41_-targeted Immuno-STAT therapy, underscoring the selectivity of this approach. However, minor changes in phenotypic markers, including increased expression of CD25 and CX3CR1, were observed in non-targeted CD8 T cells, likely reflecting bystander IL-2 signaling. Combining Immuno-STAT with PD-1 blockade augmented D^b^GP33-specific CD8 T cell responses more effectively than PD-1 blockade alone, with minor effects on the non-targeted D^b^GP276-specific population. These findings inform the clinical development of Immuno-STAT and other IL-2 therapeutics and highlight the value of coordinated TCR and IL-2 stimulation during chronic antigen exposure, alone or in combination with PD-1 blockade.

**IMPORTANCE:** Interleukin-2 (IL-2) is a key cytokine for promoting effector differentiation of antigen-specific CD8 T cells and remains an attractive agent in cancer immunotherapy, but systemic toxicity limits its clinical use. This study addresses a central challenge in IL-2-based immunotherapy: delivering IL-2 to cognate antigen-specific CD8 T cells while minimizing activation of non-targeted populations. Using a mouse model of chronic lymphocytic choriomeningitis virus (LCMV) infection, we show that the Immuno-STAT (Selective Targeting and Alteration of T cells) platform selectively expands targeted virus-specific CD8 T cells and enhances their function while limiting effects on non-targeted populations. We also show that combining Immuno-STAT with PD-1 blockade further enhances targeted virus-specific CD8 T cell responses during chronic LCMV infection. These findings provide mechanistic and preclinical support for integrating T cell receptor (TCR) specificity with IL-2 signaling to advance cancer immunotherapy and guide next-generation IL-2 therapeutics for cancer and chronic infection.

## INTRODUCTION

Interleukin-2 (IL-2), originally defined as a growth factor for T cells (1), is a critical cytokine for CD8 T cell proliferation and effector differentiation. Based on preclinical and clinical studies, human recombinant IL-2, aldesleukin (Proleukin), administered at high doses (HD), was approved by the FDA (Food and Drug Administration) in the 1990s for the treatment of metastatic melanoma and renal cancer (2). It achieved an objective response rate (ORR) of approximately 15% in these indications (2). However, several concerns related to the clinical use of IL-2, such as poor pharmacokinetics, toxicity, and activation of Foxp3^+^ regulatory CD4 T cells that constitutively express IL-2 receptor alpha (IL-2Rα or CD25), have limited its broad clinical use (3-5).

Despite these limitations, IL-2 remains one of the most attractive cytokines in cancer immunotherapy. Accordingly, significant efforts have been devoted to developing multiple IL-2-based therapeutics (3-5). One approach is to engineer IL-2R bias into IL-2 agents, such as by reducing or abolishing CD25 binding (non-α IL-2), enhancing affinity for CD122 (IL-2Rβ bias), decreasing CD132 binding (IL-2Rαβ bias), or preferentially engaging the high-affinity heterotrimeric IL-2R (IL-2Rαβγ bias). Another approach is to engineer IL-2 agents for targeted delivery to tumor sites, the tumor microenvironment (TME), or immune cells (3-5). Immune cell-targeted IL-2 therapeutics are also diverse, ranging from those targeting broad immune cell populations, including TCRβ (T cell receptor β)-expressing T cells (6) and CD8 T cells (7), to those more specifically targeting antigen-reactive cells such as PD-1-directed IL-2 (8, 9) and tumor-specific TCR-directed therapeutics (10, 11). However, it remains unclear which combinations of IL-2R bias and immune cell targeting are optimal for enhancing therapeutic efficacy while reducing toxicity. Further studies are therefore needed to characterize the *in vivo* effects of each IL-2 therapeutic on immune effector cells.

Given that immune cell-targeted IL-2 therapeutics aim to deliver IL-2 to tumor-reactive CD8 T cells while minimizing off-target toxicities, selective delivery of IL-2 signals to antigen-specific CD8 T cells may enhance their responses, promote effector differentiation, and spare non-targeted populations. One such IL-2-based drug candidate is the Immuno-STAT (Selective Targeting and Alteration of T cells) platform, which consists of a fusion protein comprising an epitope peptide, β2 microglobulin (β2M), an MHC class I allele, an affinity-attenuated IL-2 mutein, and an Fc domain. It is designed to provide TCR (T cell receptor) stimulation in conjunction with targeted IL-2 delivery to cognate epitope-specific CD8 T cells. Previous studies have shown that Immuno-STAT effectively activates and expands cognate CD8 T cells *in vitro*, in preclinical mouse tumor models, and in patients with cancer (10-13). However, the immune responses induced by Immuno-STAT remain incompletely characterized, including its effects on exhausted cognate CD8 T cells and on antigen-specific CD8 T cells not targeted by the Immuno-STAT construct.

Our recent studies have shown that IL-2 is a key cytokine that drives effector CD8 T cell differentiation during chronic LCMV infection. It acts on PD-1^+^ TCF-1^+^ stem-like subsets, also referred to as progenitor exhausted T cells (Tpex) (14-16), and modifies their differentiation trajectory, generating more effective effector cells from Tpex (17). Therefore, it is important to examine whether Immuno-STAT containing an attenuated IL-2 mutein can drive the differentiation of Tpex cells toward functionally improved effector cells as effectively as wild-type IL-2, either alone or in combination with PD-1 (programmed cell death 1) blockade. Additionally, given that human versions of Immuno-STAT constructs targeting various T cell epitopes, such as CUE-101, are currently being tested in clinical trials in patients with cancer (18), it is essential to understand how the Immuno-STAT approach influences CD8 T cell differentiation and function *in vivo*, including its effects on antigen-specific CD8 T cells not targeted by the Immuno-STAT construct in the setting of persistent antigen exposure.

Here, we studied the therapeutic effects of Immuno-STAT proteins targeting the D^b^GP_33-41_ epitope of lymphocytic choriomeningitis virus (LCMV) using a mouse model of chronic LCMV infection (14-17, 19-21). We evaluated their therapeutic potential as monotherapy and in combination with PD-1 blockade by examining their effects on antigen-specific CD8 T cell responses.

## RESULTS

### Treatment with Immuno-STAT enhances targeted D^b^GP33-specific CD8 T cell responses during chronic LCMV infection

An Immuno-STAT molecule targeting H-2D^b^-restricted LCMV GP_33-41_ epitope was developed as previously described (10, 11) (Fig. 1A). We first assessed the activity of Immuno-STAT as monotherapy during chronic LCMV infection. Chronically LCMV infected mice (> 40 days post-infection) were either left untreated or treated with Immuno-STAT (15 mg/kg intraperitoneally (i.p.)) twice daily for 7-10 days, and LCMV-specific, both Immuno-STAT targeted D^b^GP33-specific and non-targeted D^b^GP276-specific, CD8 T cell responses were analyzed in spleen, liver, and blood (Fig. 1B). The number of Immuno-STAT-targeted D^b^GP33-specific CD8 T cells assessed by tetramer staining was increased in all tissues examined (Fig. 1C and D), while non-targeted D^b^GP276-specific CD8 T cells were largely unaffected with only a modest increase observed in the blood (Fig. 1E and F). In addition, expanded Immuno-STAT-targeted D^b^GP33-specific CD8 T cells were functional in terms of producing interferon gamma (IFN-γ) effector cytokine after *ex vivo* stimulation with cognate peptide (Fig. 2A), while D^b^GP276-specific CD8 T cells did not show improved effector cytokine production (Fig. 2B). These results demonstrate that the Immuno-STAT platform selectively enhances epitope-specific CD8 T cell responses during chronic LCMV infection.

**Figure 1.**
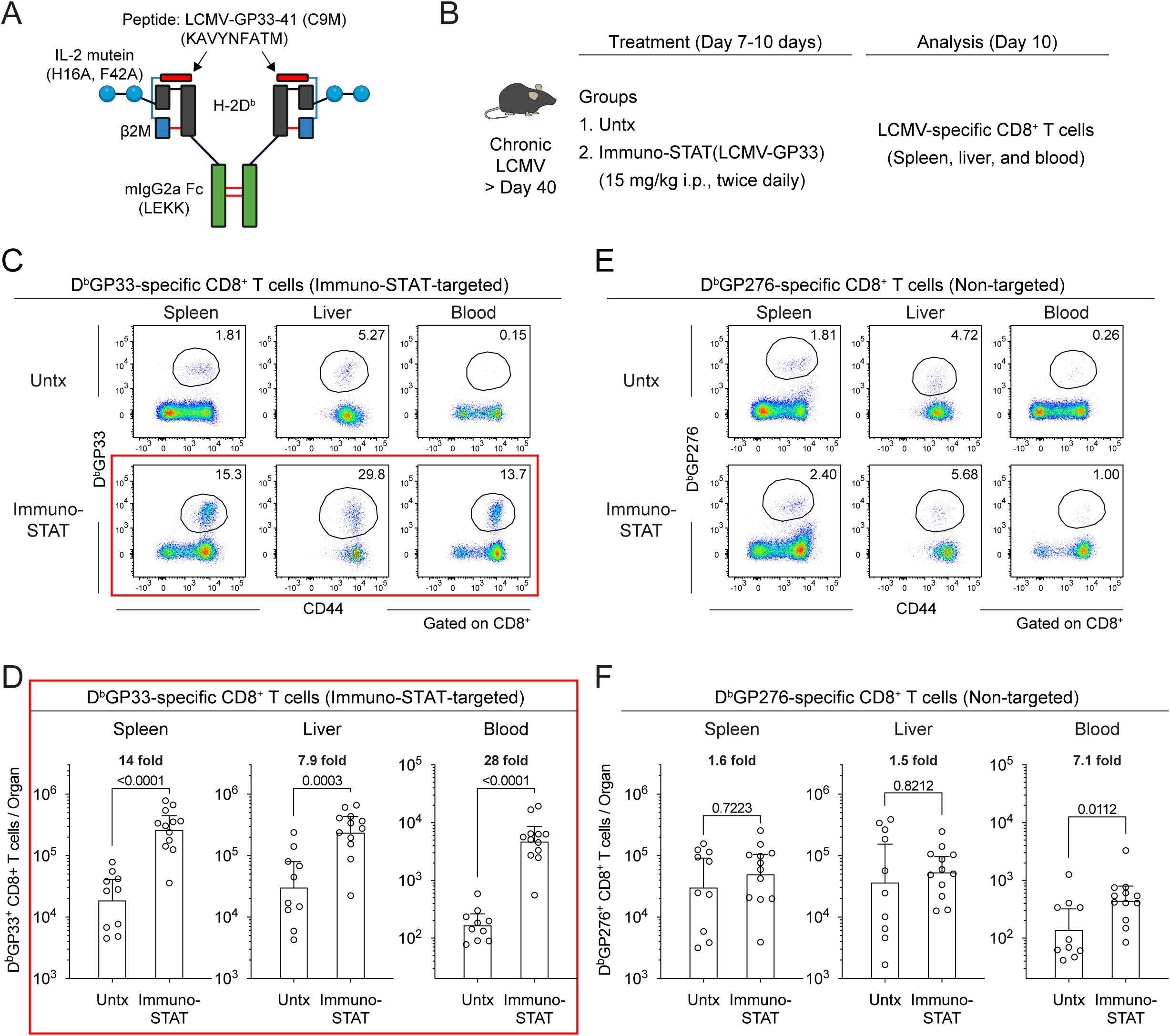
Immuno-STAT selectively enhances targeted D^b^GP33-specific CD8 T cell responses during chronic LCMV infection. (A) Immuno-STAT design. Immuno-STAT proteins consisting of H-2D^b^ loaded with the LCMV GP33-41 peptide (KAVYNFATM) were genetically fused to a human IL-2 mutein containing the H16A and F42A point mutations and to an effector-attenuated murine IgG2a Fc domain. (B) Experimental design for panels (C-F). Chronically LCMV-infected mice (> 40 days post-infection) were left untreated or treated with Immuno-STAT (15 mg/kg, i.p., twice daily) for 7-10 days, followed by analysis of LCMV-specific CD8 T cells in the spleen, liver, and blood. (C) Representative flow cytometry plots showing tetramer staining of Immuno-STAT-targeted D^b^GP33-specific CD8 T cells in the indicated tissues. (D) Summary plots showing the number of D^b^GP33-specific CD8 T cells. (E) Representative flow cytometry plots showing tetramer staining of non-targeted D^b^GP276-specific CD8 T cells in the indicated tissues. (F) Summary plots showing the number of D^b^GP276-specific CD8 T cells. Results were pooled from 3 experiments with n=3-4 mice per group in each experiment. Statistical comparisons were performed using the unpaired Mann-Whitney test (D, F). Immuno-STAT-targeted D^b^GP33-specific CD8 T cells are highlighted by red boxes (C and D). Bars and error bars represent the geometric mean and 95% confidence interval (D, F). Untx, untreated.

**Figure 2.**
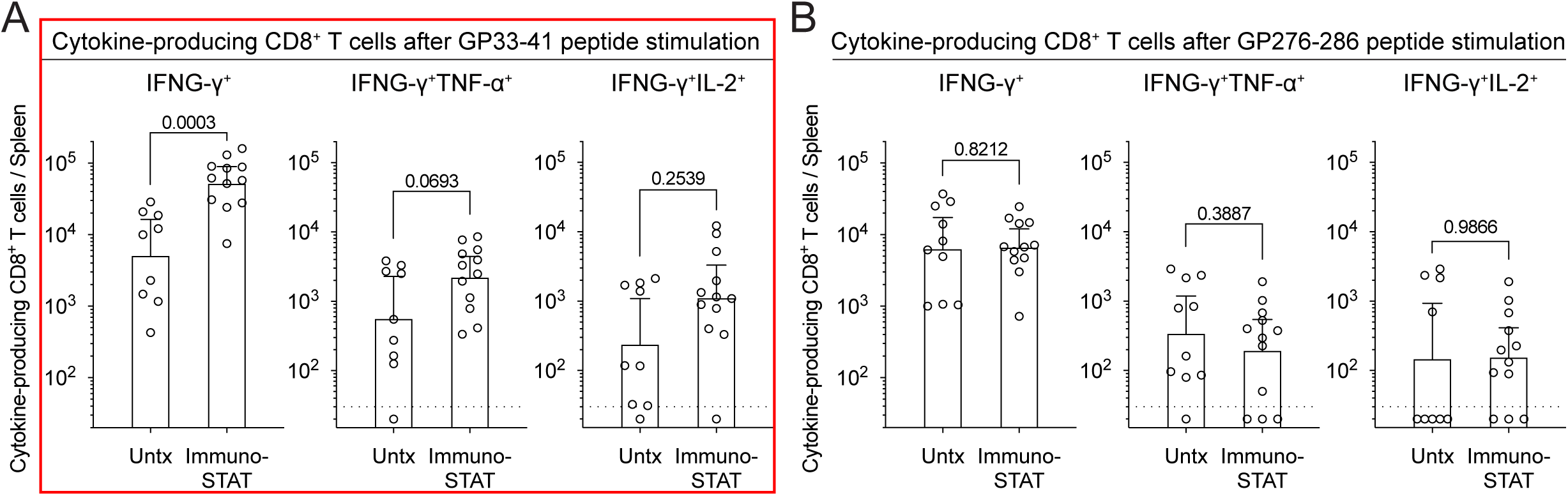
Immuno-STAT selectively augments effector cytokine production in targeted D^b^GP33-specific CD8 T cells during chronic LCMV infection. Chronically LCMV-infected mice (> 40 days post-infection) were left untreated or treated with Immuno-STAT (15 mg/kg, i.p., twice daily) for 7-10 days. (A, B) Splenocytes isolated from each treatment group were stimulated with GP33-41 (A) or GP276-286 peptide (B) for 5 hours, followed by analysis of cytokine-producing CD8 T cells. Results were pooled from 3 experiments with n=3-4 mice per group in each experiment. Statistical comparisons in panels A and B were performed using the unpaired Mann-Whitney test. Immuno-STAT-targeted D^b^GP33-specific CD8 T cell responses are highlighted by a red box (A). Bars and error bars represent the geometric mean and 95% confidence interval (A, B). Untx, untreated.

### Immuno-STAT selectively delivers IL-2 signals to targeted D^b^GP33-specific CD8 T cells and promotes effector differentiation during chronic LCMV infection

IL-2 is a key cytokine to drive effector CD8 T cell differentiation during chronic LCMV infection, acting on PD-1^+^ TCF-1^+^ stem-like CD8 T cells and generating better effectors similar to those seen after an acute LCMV infection (17). To address if Immuno-STAT mediated IL-2 signal delivery effectively changes CD8 T cell differentiation trajectory, we characterized the phenotype of Immuno-STAT-targeted D^b^GP33-specific and non-targeted D^b^GP276-specific CD8 T cells by multicolor flow cytometry after Immuno-STAT monotherapy during chronic LCMV infection. Immuno-STAT therapy promoted the proliferation and effector differentiation of D^b^GP33-specific CD8 T cells based on increased expression of Ki-67, granzyme B, CX3CR1, CD218a, and CD25 as well as decreased expression of CD101 compared to untreated mice (Fig. 3A). In contrast, non-targeted D^b^GP276-specific CD8 T cells in mice treated with Immuno-STAT did not show increased expression of Ki-67 and granzyme B compared to those in untreated mice (Fig. 3B), indicating only modest changes in their proliferation and effector differentiation. In contrast, CD25 expression was upregulated by Immuno-STAT therapy in both targeted D^b^GP33-specific and non-targeted D^b^GP276-specific CD8 T cells, implying some modest off-target delivery of IL-2 signals to non-targeted T cells under this treatment regimen (Fig. 3A and B). Consistent with this, increased expression of CX3CR1, suggesting effector differentiation, was also observed in non-targeted D^b^GP276-specific CD8 T cells relative to untreated mice (Fig. 3B). Similarly, there was a trend of increased and decreased expression of CD218a and CD101 in non-targeted D^b^GP276-specific CD8 T cells, respectively, although these changes did not reach statistical significance (Fig. 3C). The observed changes in these phenotypic markers clearly indicate that Immuno-STAT therapy selectively delivered TCR and IL-2 signals to D^b^GP33-specific CD8 T cells, promoting their proliferation and effector differentiation, while modest off-target IL-2 signals were received by non-targeted D^b^GP276-specific CD8 T cells during chronic LCMV infection (Fig. 3C). Taken together, these results further support that Immuno-STAT molecules act selectively on targeted CD8 T cell population and promote their effector differentiation. Single-agent Immuno-STAT therapy did not improve viral control in chronic LCMV infection (mean spleen PFU/g: untreated, 2.6 x 10^6^; treated, 1.4 x 10^7^). This outcome was not unexpected, given that the chronic LCMV model is a stringent system marked by profound T cell exhaustion and lifelong viremia, and targeting a single epitope with Immuno-STAT may be insufficient to control systemic viral dissemination.

**Figure 3.**
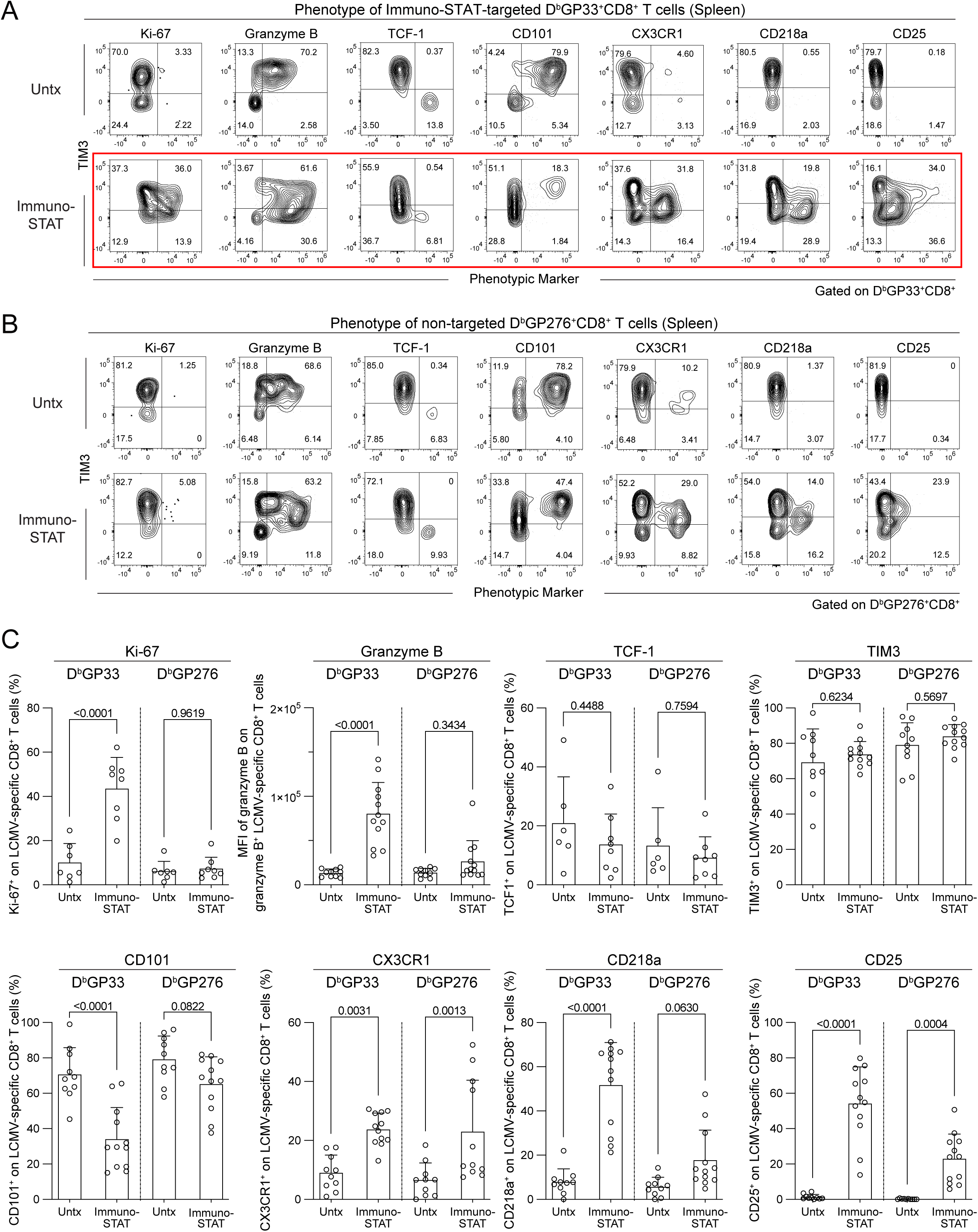
Effects of Immuno-STAT on the phenotype of targeted and non-targeted LCMV-specific CD8 T cells after Immuno-STAT treatment during chronic infection. Chronically LCMV-infected mice (> 40 days post-infection) were left untreated or treated with Immuno-STAT for 7-10 days, followed by phenotypic analysis of Immuno-STAT-targeted D^b^GP33- and non-targeted D^b^GP276-specific CD8 T cells in the spleen. (A, B) Representative flow cytometry plots showing co-expression of TIM3 and the indicated phenotypic markers on D^b^GP33-specific CD8 T cells (A) and D^b^GP276-specific CD8 T cells (B). (C) Summary plots showing expression of various phenotypic markers on D^b^GP33- and D^b^GP276-specific CD8 T cells. Results were pooled from 2-3 experiments with n=3-4 mice per group in each experiment. Statistical comparisons in (C) were performed using two-way ANOVA with Šídák’s correction for multiple comparisons. A red box highlights the phenotypic changes observed in Immuno-STAT-treated DbGP33-specific CD8 T cells (A). Bars and error bars represent the mean and standard deviation (C). Untx, untreated.

### Immuno-STAT therapy in combination with PD-1 blockade further expands targeted D^b^GP33-specific CD8 T cells during chronic LCMV infection

Given the promising results of Immuno-STAT monotherapy in enhancing targeted CD8 T cell responses, we next examined effects of Immuno-STAT therapy in combination with PD-1 blockade. Chronically LCMV infected mice (> 40 days post-infection) were either left untreated or treated with anti-PD-L1 antibody alone (200 μg i.p., every 3 days), Immuno-STAT alone, or the combination of anti-PD-L1 antibody and Immuno-STAT for 10 days, followed by the analysis of LCMV-specific D^b^GP33^+^ CD8 T cells in spleen, liver, and blood (Fig. 4A). Addition of PD-1-therapy to Immuno-STAT was beneficial to enhance Immuno-STAT-targeted D^b^GP33-specific CD8 T cell responses in all the tissues examined (Fig. 4B and C). Cytokine-producing cells show similar trends, with combination therapy effectively enhancing the functionality of targeted D^b^GP33-specific CD8 T cells relative to either monotherapy (Fig. 5A and B).

**Figure 4.**
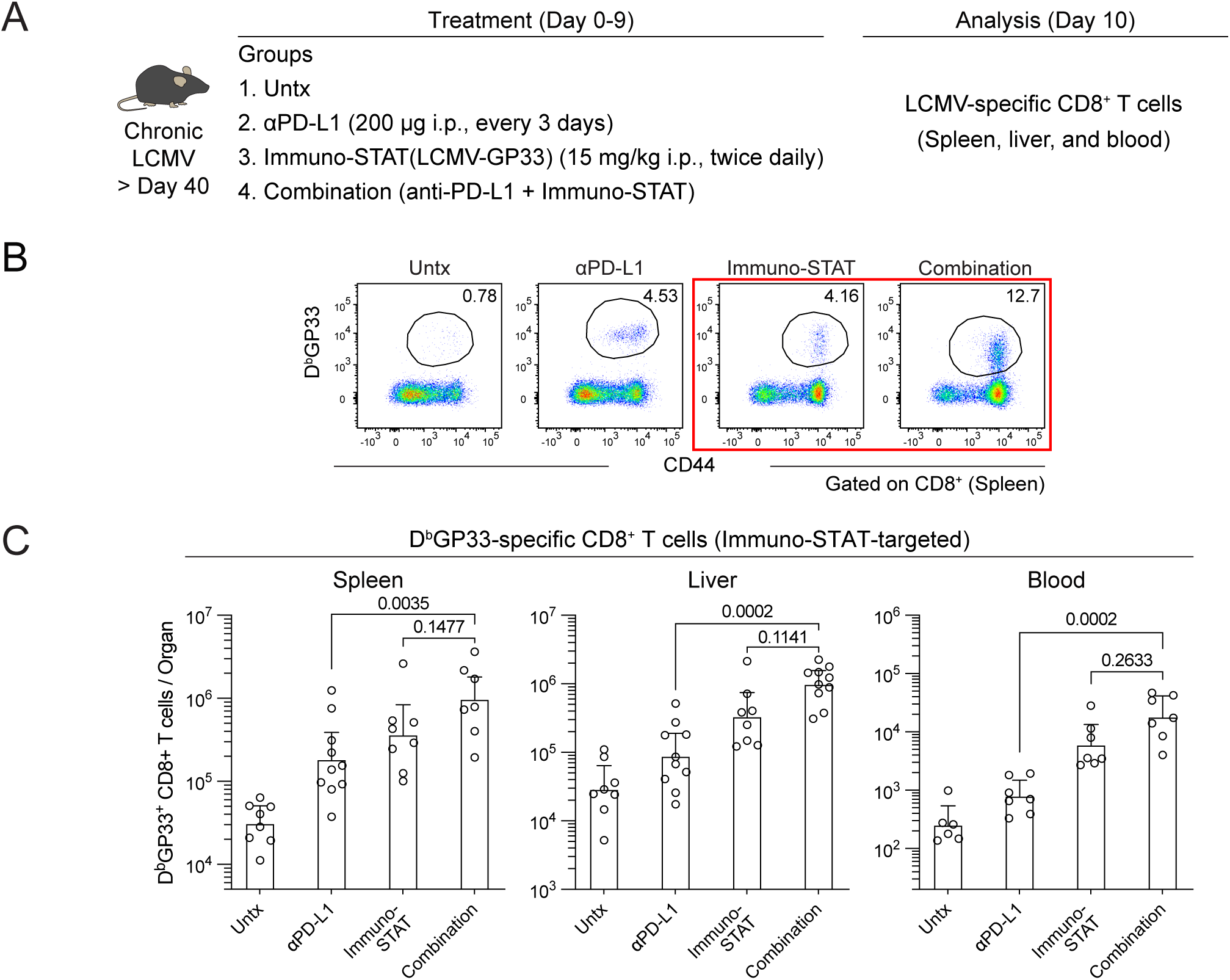
Combination treatment with Immuno-STAT and anti-PD-L1 further enhances targeted D^b^GP33-specific CD8 T cell responses during chronic LCMV infection. (A) Experimental design. Chronically LCMV-infected mice (> 40 days post-infection) were left untreated or treated with anti-PD-L1 antibody alone, Immuno-STAT alone, or a combination of anti-PD-L1 antibody and Immuno-STAT for 10 days, followed by analysis of LCMV-specific CD8 T cells. (B) Representative flow cytometry plots showing tetramer staining of Immuno-STAT-targeted D^b^GP33-specific CD8 T cells in the spleen. (C) Summary plots showing the number of D^b^GP33-specific CD8 T cells. Results were pooled from 2-3 experiments with n=1-4 mice per group in each experiment. Statistical comparisons in (C) were performed using the Kruskal-Wallis test with Dunn’s correction. Immuno-STAT-targeted D^b^GP33-specific CD8 T cell responses are highlighted by a red box (B). Bars and error bars represent the geometric mean and 95% confidence interval (C). Untx, untreated.

**Figure 5.**
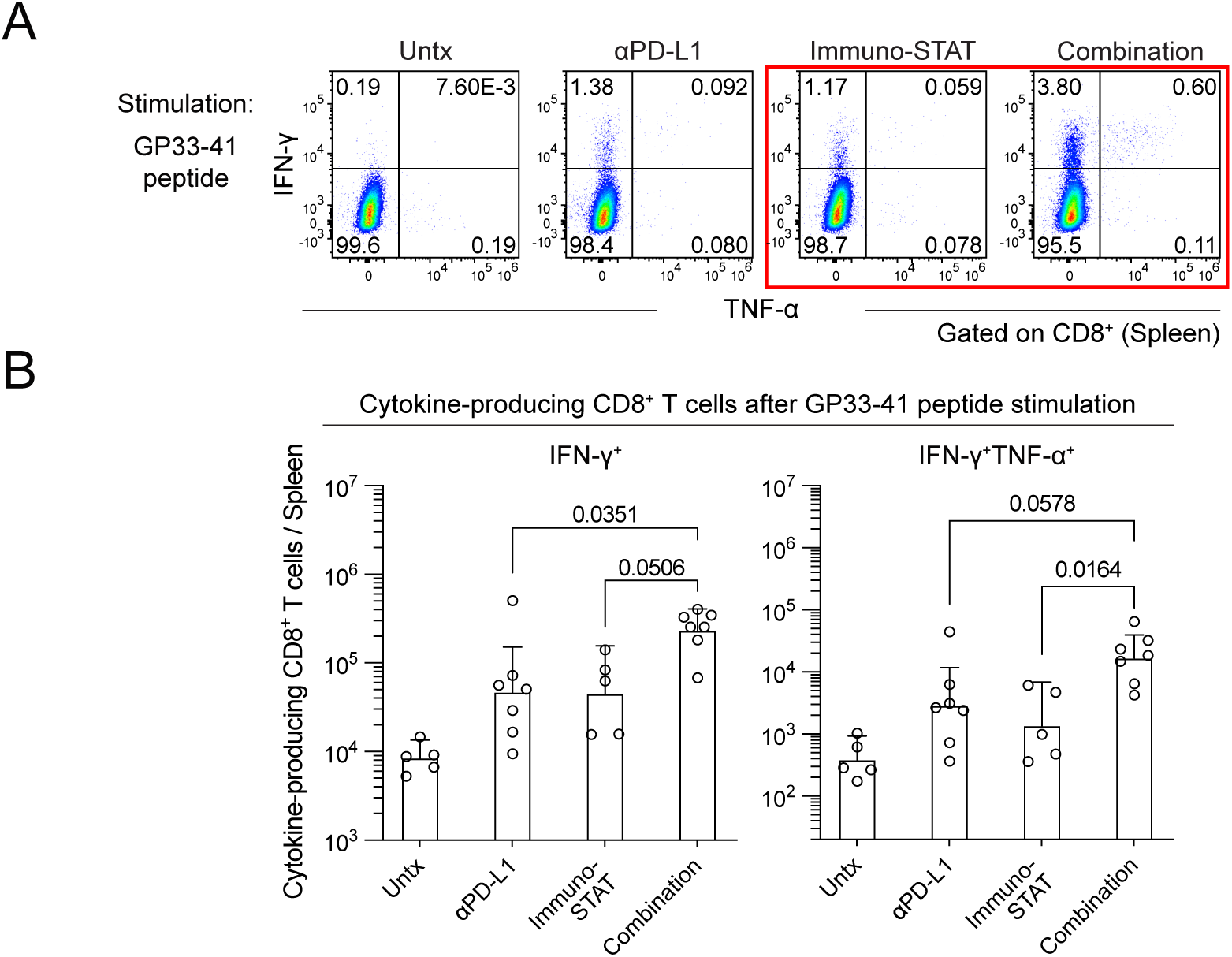
Combining Immuno-STAT therapy with PD-1 blockade further enhances effector cytokine production in targeted D^b^GP33-specific CD8 T cells during chronic LCMV infection. Chronically LCMV-infected mice (> 40 days post-infection) were left untreated or treated with anti-PD-L1 antibody alone, Immuno-STAT alone, or a combination of anti-PD-L1 antibody and Immuno-STAT for 10 days. Splenocytes from each treatment group were stimulated with GP33-41 peptide for 5 hours, followed by analysis of cytokine-producing CD8 T cells. (A) Representative flow cytometry plots for the expression of IFN-γ and TNF-α in the indicated groups. (B) Summary plots showing the number of cytokine-producing CD8 T cells. Results were pooled from 2 experiments with n=1-4 mice per group in each experiment. Statistical comparisons in (B) were performed using the Kruskal-Wallis test with Dunn’s correction. A red box highlights cytokine-producing Immuno-STAT-targeted D^b^GP33-specific CD8 T cells (A). Bars and error bars represent the geometric mean and 95% confidence interval (B). Untx, untreated.

### Immuno-STAT alone or in combination with PD-1 blockade drives effector differentiation in D^b^GP33-specific CD8 T cells during chronic LCMV infection

Our recent studies show that IL-2 signals, when optimally delivered to antigen-specific CD8 T cells, result in the generation of better effector cells with enhanced therapeutic potential from the PD-1^+^ TCF-1^+^ stem-like subset in chronic LCMV infection and mouse cancer models (8, 17). To determine whether Immuno-STAT induces similar effects in targeted CD8 T cells, we conducted detailed phenotypic characterization of D^b^GP33-specific and D^b^GP276-specific CD8 T cells by multicolor spectral flow cytometry and high-dimensional analysis with UMAP projection and FlowSOM clustering. Concatenated samples of both D^b^GP33-specific CD8 T cells across all treatment groups were comprised of 3 clusters (Fig. 6A and B). All subsets from LCMV-specific CD8 T cells express high levels of PD-1 and TOX compared to naïve CD8 T cells (Fig. S1) (22-24), while cells in cluster 1 were uniquely discriminated by TCF-1 expression, indicating their identity as a stem-like subset (14-16). Cells in cluster 1 also express SLAMF6 and CD73 compared to the other two subsets, but they did not express effector molecules of granzyme B and were negative for the inhibitory receptors CD101 and TIM3 (Fig. 6C). Cells in cluster 2 uniquely expressed Ki-67, and showed higher levels of granzyme B, cytokine receptors including CD25, CD218a, and CD119, the transcription factor T-bet, and migration markers including Ly6C, and CX3CR1, but were negative for TCF-1 and CD101 and express lower levels of CD69 and TIM3 (Fig. 6C). These characterizations were consistent with cluster 2 being represented by an effector population (20, 21). Cells in cluster 3 show the classical exhausted phenotype, characterized by non-dividing (Ki-67 negative), expression of high levels of CD101 and TIM3, but still retaining some levels of the effector molecule granzyme B (Fig. 6C). The exhausted cells in cluster 3 also exhibited lower expression of inflammatory cytokine receptors such as CD25, CD218a, and CD119; migration markers including CD44, Ly-6C, and CX3CR1; and the transcription factor T-bet, while re-expressing the tissue-residency marker CD69 and expressing higher levels of the inhibitory receptors CD101 and TIM3 than the other two T cell subsets, in line with previous studies (Fig. 6C) (16, 20, 25). In untreated mice with chronic LCMV, the LCMV-specific D^b^GP33^+^ CD8 T cell population comprised 17.6% ± 12.9% stem-like cells (cluster 1), 8.1% ± 7.2% effector cells (cluster 2), and 74.3% ± 14.8% exhausted cells (cluster 3) (mean ± SD; Fig. 6D). PD-1 blockade increased the proportion of cluster 2 effector cells, although this change was not statistically significant, consistent with a proliferative burst from stem-like CD8 T cells induced by PD-1 blockade. The proportion of cluster 1 cells was slightly decreased but remained detectable, suggesting self-renewal within this subset (Fig. 6D). In contrast, single-agent Immuno-STAT treatment significantly increased the proportion of cluster 2 effector cells among Immuno-STAT-targeted D^b^GP33-specific CD8 T cells to approximately 80% (Fig. 6D). Combination therapy similarly promoted effector differentiation of Immuno-STAT-targeted D^b^GP33-specific CD8 T cells (Fig. 6D). However, adding Immuno-STAT to anti-PD-L1 treatment did not improve viral control during chronic LCMV infection (mean spleen PFU/g: untreated, 2.6 x 10^6^; anti-PD-L1 treated, 1.1 x 10^6^; Immuno-STAT treated, 1.2 x 10^7^; combination treatment, 9.5 x 10^6^). These findings underscore the stringency of this lifelong viremic model of T cell exhaustion and suggest that targeting a single epitope with Immuno-STAT is insufficient to control systemic viral spread.

**Figure 6.**
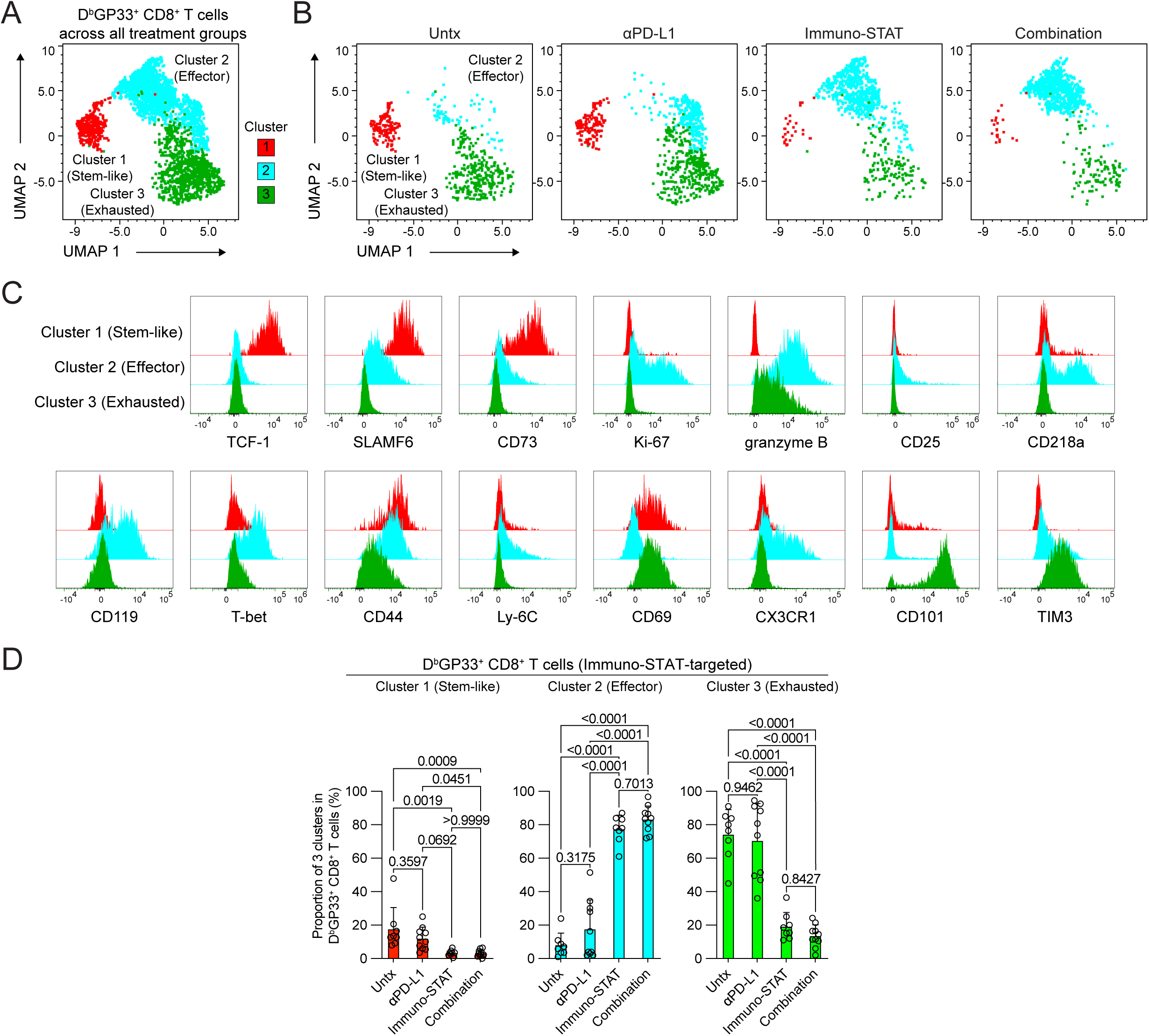
Immuno-STAT alone or in combination with anti-PD-L1 enhances the generation of effector cells in targeted D^b^GP33-specific CD8 T cell populations during chronic LCMV infection. Chronically LCMV-infected mice (> 40 days post-infection) were left untreated or treated with anti-PD-L1 antibody alone, Immuno-STAT alone, or a combination of anti-PD-L1 antibody and Immuno-STAT for 10 days, followed by analysis of LCMV-specific CD8 T cells. (A, B) Representative UMAP projections with FlowSOM clustering showing three clusters of concatenated D^b^GP33-specific CD8 T cells isolated from spleens across the four treatment groups (A) and the corresponding cluster distribution in each treatment group (B). (C) Representative histograms showing expression of the indicated phenotypic markers on D^b^GP33-specific CD8 T cells within each of the three clusters. (D) Proportions of the three D^b^GP33-specific CD8 T cell clusters in the different treatment groups. Results were pooled from 2-3 experiments with n=1-4 mice per group in each experiment. Statistical comparisons were performed using one-way ANOVA with Tukey’s correction for multiple comparisons (D). Bars and error bars represent the mean and standard deviation (D). Untx, untreated.

### Immuno-STAT in combination with PD-1 therapy selectively enhances targeted D^b^GP33-specific CD8 T cell responses and improves viral control in the chronic LCMV infection model with CD4 T cell help

The results presented thus far were obtained in a stringent model of chronic LCMV infection characterized by lifelong viremia in the absence of LCMV-specific CD4 T cells (19, 26). We therefore investigated the effects of Immuno-STAT alone and in combination with PD-1 blockade in a less stringent model of chronic LCMV infection, in which mice were infected with LCMV clone 13 without transient CD4 T cell depletion (19). This less stringent model allows assessment of treatment effects on LCMV-specific CD8 T cells in the presence of LCMV-specific CD4 T cells (17). At approximately 3 weeks post-infection, mice were left untreated or treated with anti-PD-L1 antibody alone, Immuno-STAT alone, or the combination of Immuno-STAT and anti-PD-L1 antibody for 7-10 days, followed by analysis of LCMV-specific CD8 T cells and viral titers in spleen and liver (Fig. 7A). As in the more stringent model, addition of Immuno-STAT to PD-1 therapy further enhanced Immuno-STAT-targeted D^b^GP33-specific CD8 T cell responses, as measured by D^b^GP33 tetramer staining, in spleen, liver, and blood compared with PD-1 monotherapy, although Immuno-STAT alone was largely sufficient to increase the number of these cells (Fig. 7B and C). In contrast, non-targeted D^b^GP276-specific CD8 T cell responses were not enhanced by Immuno-STAT compared with untreated mice, and combination therapy did not further improve D^b^GP276-specific CD8 T cell responses beyond PD-1 monotherapy (Fig. 7B and D). Similar results were observed for effector cytokine-producing cells after *ex vivo* stimulation of splenocytes with GP33-41 or GP276-286 peptide (Fig. 8A and B). Additionally, Immuno-STAT selectively promoted the proliferation and effector differentiation of D^b^GP33-specific CD8 T cells (Fig. 9A), while modest off-target effects were also observed in non-targeted D^b^GP276-specific CD8 T cells, including changes in the expression of CD25, CD218a, and CD101 after Immuno-STAT monotherapy or combination therapy (Fig. 9B). Interestingly, combination therapy with Immuno-STAT and PD-1 blockade was associated with improved viral control in the spleen and liver in this less stringent model of chronic LCMV infection (Fig. S2). Of note, PD-1 blockade was required to achieve improved viral control, and enhanced T cell responses alone were not sufficient in this chronic LCMV model, as previously described (17), highlighting a potential limitation of targeting a single epitope in LCMV infection.

**Figure 7.**
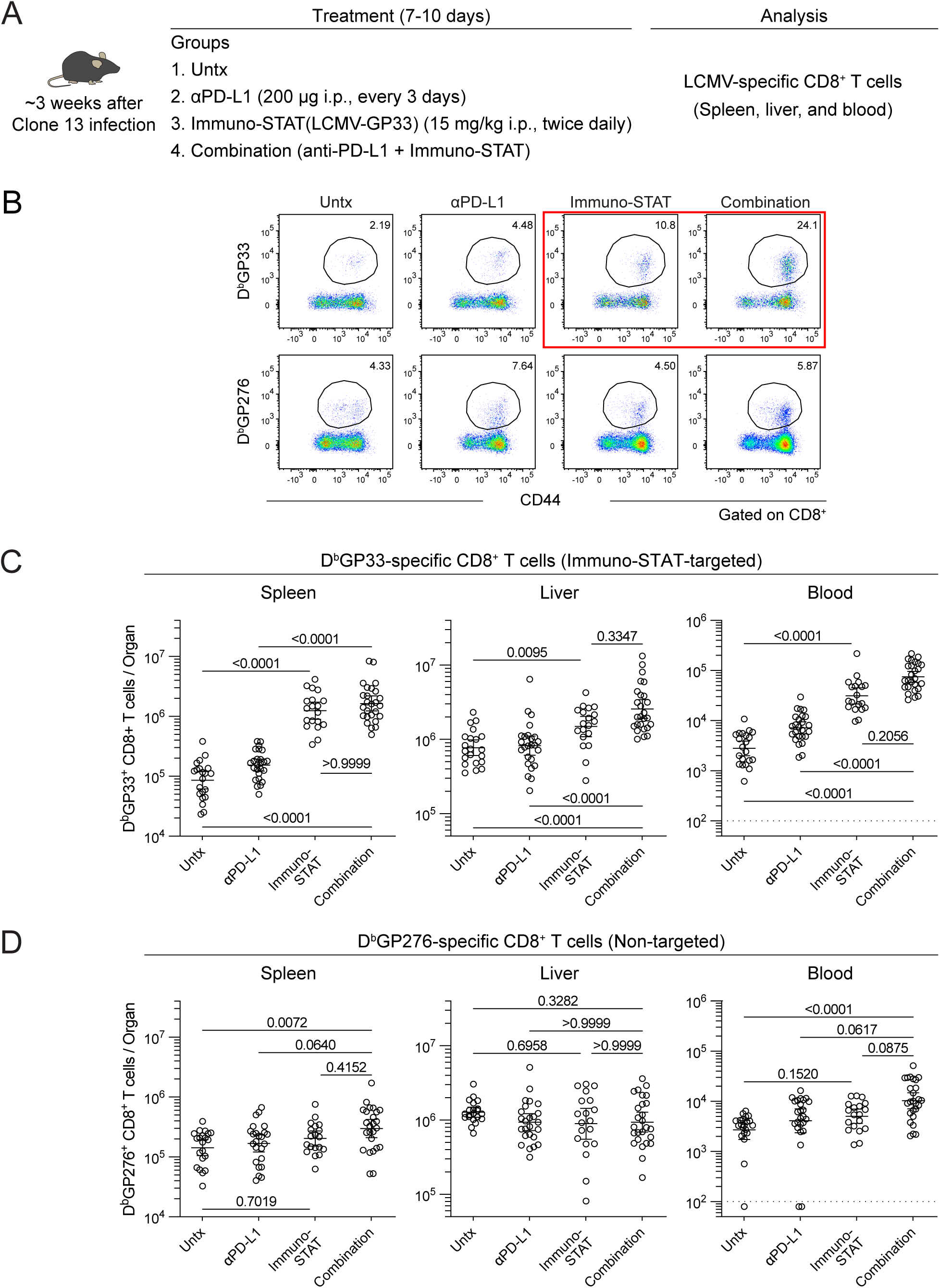
Targeted and non-targeted virus-specific CD8 T cell responses after Immuno-STAT therapy alone or in combination with anti-PD-L1 treatment in a chronic LCMV model with CD4 T cell help. (A) Experimental design. Chronically LCMV-infected mice (19-25 days post-infection) were left untreated or treated with anti-PD-L1 antibody alone, Immuno-STAT alone, or a combination of anti-PD-L1 antibody and Immuno-STAT for 7-10 days, followed by analysis of LCMV-specific CD8 T cell responses. (B) Representative flow cytometry plots showing tetramer staining of Immuno-STAT-targeted D^b^GP33-specific and non-targeted D^b^GP276-specific CD8 T cells in the spleen. (C, D) Summary plots showing the number of Immuno-STAT-targeted D^b^GP33-specific (C) and non-targeted D^b^GP276-specific CD8 T cells (D) in the indicated tissues. Results were pooled from 5-7 experiments with n=1-5 mice per group in each experiment. Statistical comparisons were performed using the Kruskal-Wallis test with Dunn’s correction for multiple comparisons (C, D). A red box in panel B highlights Immuno-STAT-targeted D^b^GP33-specific CD8 T cells. Bars and error bars represent the geometric mean and 95% confidence interval (C, D). Untx, untreated.

**Figure 8.**
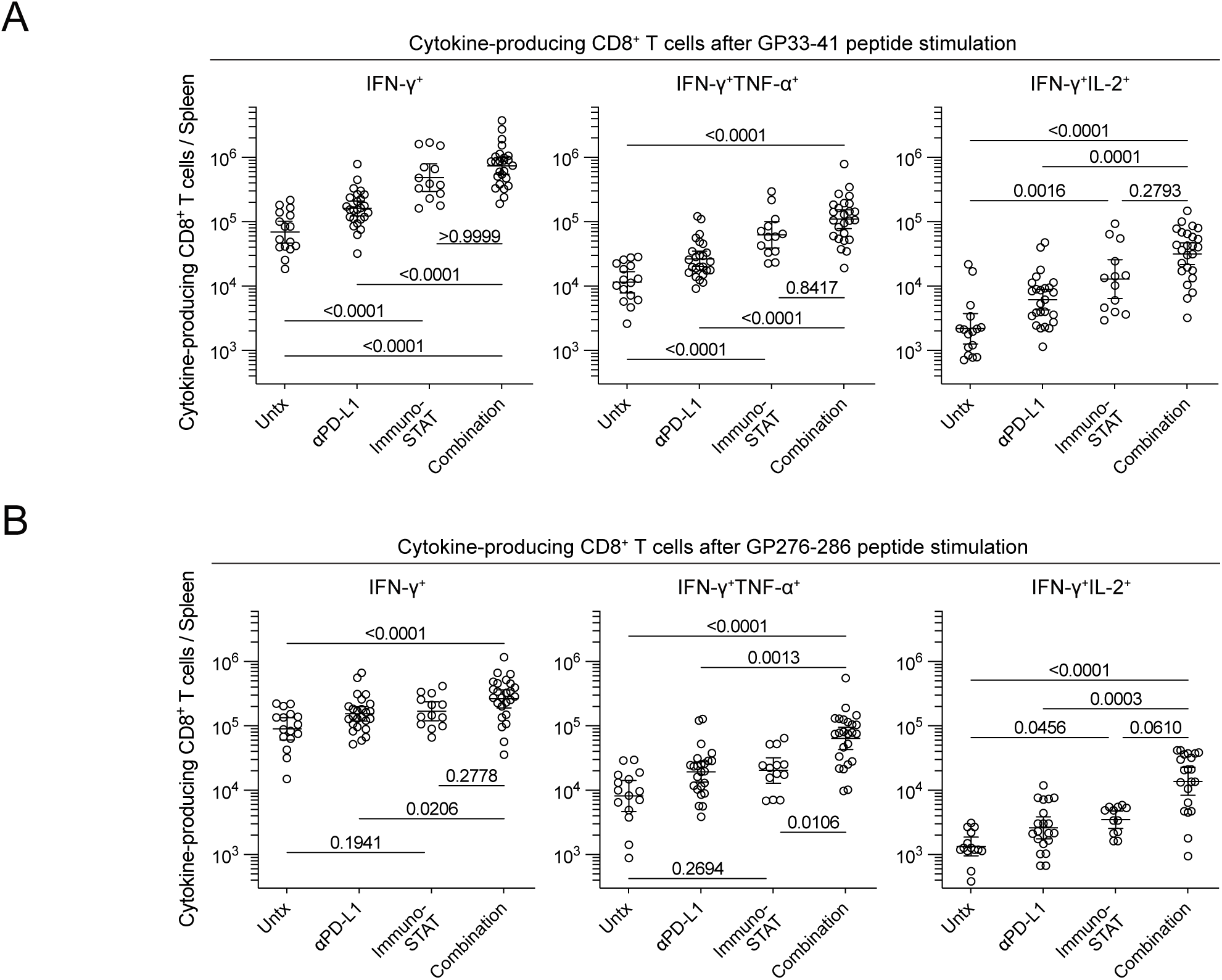
Cytokine production by targeted and non-targeted virus-specific CD8 T cells after Immuno-STAT therapy alone or in combination with anti-PD-L1 treatment in a chronic LCMV model with CD4 T cell help. Chronically LCMV-infected mice (19-25 days post-infection) were left untreated or treated with anti-PD-L1 antibody alone, Immuno-STAT alone, or a combination of anti-PD-L1 antibody and Immuno-STAT for 7-10 days. Splenocytes from each treatment group were stimulated with GP33-41 or GP276-286 peptide for 5 hours, followed by analysis of cytokine-producing CD8 T cells. (A, B) Summary plots showing the number of cytokine-producing CD8 T cells in response to GP33-41 peptide (A) or GP276-286 peptide (B). Results were pooled from 7 experiments with n=1-5 mice per group in each experiment. Statistical comparisons were performed using the Kruskal-Wallis test with Dunn’s correction for multiple comparisons (A, B). Bars and error bars represent the geometric mean and 95% confidence interval (A, B). Untx, untreated.

**Figure 9.**
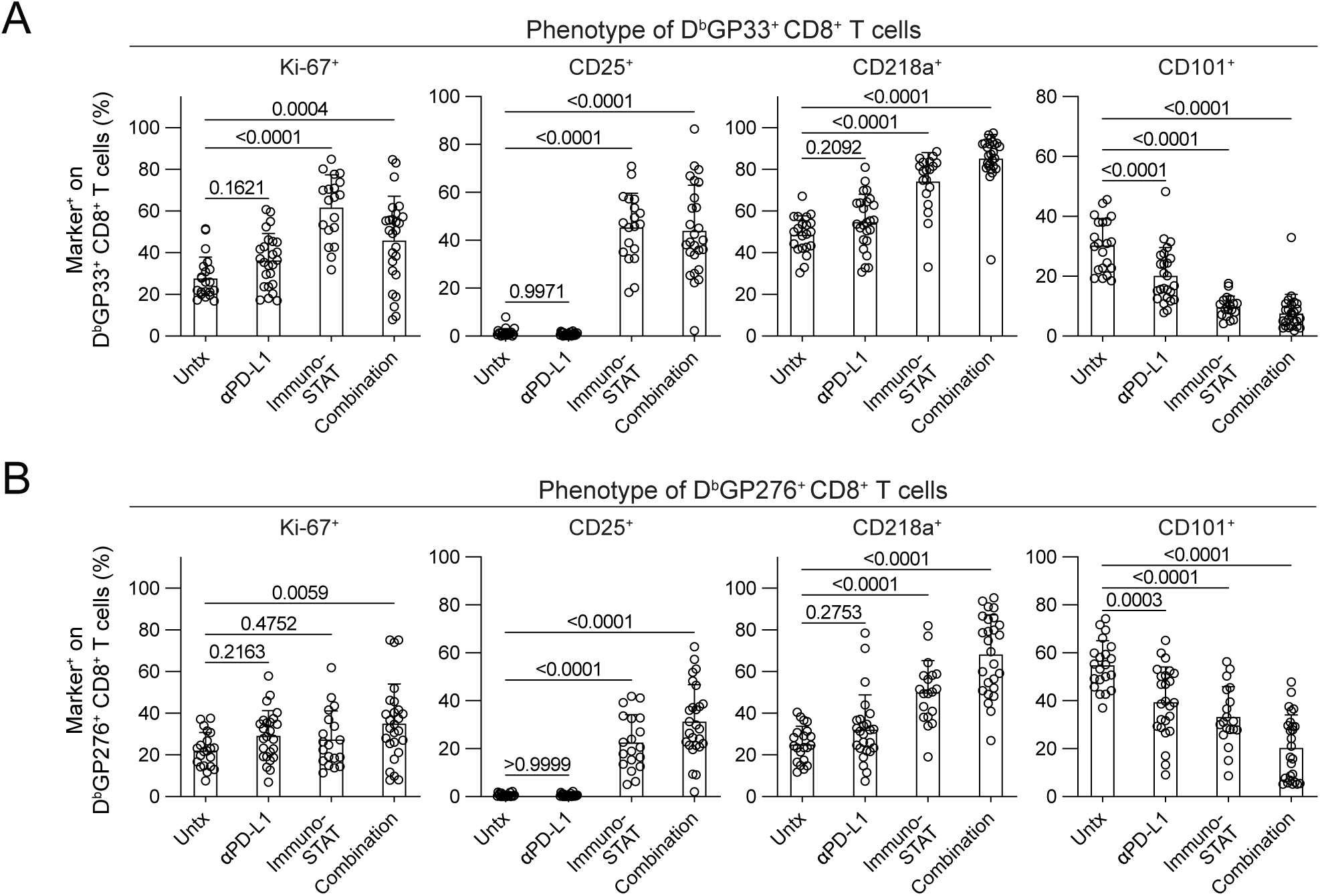
Phenotypic characterization of targeted and non-targeted virus-specific CD8 T cells after Immuno-STAT therapy alone or in combination with anti-PD-L1 treatment in a chronic LCMV model with CD4 T cell help. Chronically LCMV-infected mice (19-25 days post-infection) were left untreated or treated with anti-PD-L1 antibody alone, Immuno-STAT alone, or a combination of anti-PD-L1 antibody and Immuno-STAT for 7-10 days. (A, B) Summary plots showing the expression of various markers on Immuno-STAT-targeted D^b^GP33-specific CD8 T cells (A) and non-targeted D^b^GP276-specific CD8 T cells (B). Results were pooled from 5-7 experiments with n=1-5 mice per group in each experiment. Statistical comparisons were performed using one-way ANOVA with Tukey’s correction for multiple comparisons (A, B). Bars and error bars represent the mean and standard deviation (A, B). Untx, untreated.

## DISCUSSION

Immuno-STAT is a novel therapeutic platform designed to enable selective TCR stimulation and targeted delivery of affinity-attenuated IL-2 to antigen-specific CD8 T cells (10, 11, 18). Using mouse models of chronic LCMV infection, well-defined systems for studying T cell responses under persistent antigenic stimulation (14-17, 19-21), we found that Immuno-STAT, a murine surrogate of the CUE-100 molecule, selectively enhanced targeted LCMV D^b^GP33-specific CD8 T cell responses and promoted IL-2-driven effector differentiation, both as monotherapy and in combination with PD-1 blockade. These findings support the idea that TCR-targeted IL-2 therapeutics represent a promising approach for selective mobilization and activation of antigen-specific CD8 T cells.

The Immuno-STAT-driven effector differentiation observed in targeted D^b^GP33-specific CD8 T cell responses was consistent with our previous studies using wild-type IL-2, in which combination therapy with IL-2 and PD-1 blockade synergistically enhanced virus-specific CD8 T cell responses in the chronic LCMV model (17). This combination therapy targeted PD-1^+^ TCF-1^+^ stem-like CD8 T cells, promoted their proliferation and differentiation, and generated transcriptionally and epigenetically distinct effector CD8 T cells during chronic LCMV infection (17). These IL-2-driven effector CD8 T cells resembled highly functional effectors from acute viral infections, overcoming epigenetic barriers associated with exhausted CD8 T cells and enabling superior viral control (17). Consistent with those earlier studies, Immuno-STAT selectively increased proliferation of targeted D^b^GP33-specific CD8 T cells and promoted their differentiation toward an effector state, reflected by an increased frequency of cells with an effector phenotype (cluster 2 in Fig. 6).

This effector cluster was characterized by upregulation of the effector molecule granzyme B, inflammatory cytokine receptors (CD25, CD119, and CD218a), molecules related to migration and adhesion (CX3CR1 and Ly-6C), and the transcription factor T-bet, along with reduced expression of inhibitory receptors such as CD101 and TIM3. This phenotype was consistent with our previous characterization of IL-2-driven effector CD8 T cells during chronic LCMV infection (17). Together, these results indicate that Immuno-STAT efficiently engaged the targeted D^b^GP33-specific CD8 T cell population, acted on the PD-1^+^ TCF-1^+^ stem-like subset, and promoted its proliferation and effector differentiation. However, we did not confirm the lineage relationship among the stem-like, effector, and exhausted subsets in the present study. Transcriptional and epigenetic profiling of D^b^GP33-specific CD8 T cells is also warranted in future studies.

Whereas our previous studies demonstrated that wild-type IL-2 elicited greater therapeutic effects than an attenuated IL-2 mutein through high-affinity heterotrimeric IL-2Rs (CD25, CD122, and CD132) upregulated on LCMV-specific CD8 T cells (17), Immuno-STAT exerts its therapeutic effects through a different mechanism involving TCR stimulation and targeted delivery of an attenuated IL-2 mutein. This mechanism is enabled by the design of Immuno-STAT proteins, which are fusion proteins containing an epitope peptide, β2M, an MHC class I allele, an affinity-attenuated IL-2 mutein, and the Fc domain, enabling targeted delivery to CD8 T cells via their cognate antigen-specific TCRs (10, 11). The IL-2 mutein of Immuno-STAT contains F42A and H16A mutations in the human IL-2 sequence, which reduce affinity for human CD25 (110-fold reduction) and CD122 (3-fold reduction), respectively (10). Correspondingly, Immuno-STAT with this IL-2 mutein showed reduced (∼2600-fold) functional activity in stimulating the mouse CTLL-2 cell line, which lacks the target TCR, compared with human wild-type IL-2 (10). The unique design thus enables Immuno-STAT to selectively target cells via TCR engagement while reducing conventional IL-2R-mediated effects through the attenuated IL-2 mutein.

Our results demonstrated that Immuno-STAT at the dosages administered in this preclinical study exhibited modest off-target effects on non-targeted LCMV-specific CD8 T cells, especially when combined with PD-1 blockade during chronic LCMV infection. Non-targeted D^b^GP276-specific CD8 T cells showed slight enhancement of their effector differentiation, likely due to IL-2 mutein binding to high-affinity heterotrimeric IL-2R, which are formed through the upregulation of CD25 on these cells due to persistent viral antigenic stimulations. Nonetheless, these off-target IL-2 effects on D^b^GP276-specific CD8 T cells were not sufficient for their significant expansion. These findings suggest that chronic antigenic stimulation creates a favorable environment for antigen-specific CD8 T cells to capture IL-2 despite affinity attenuation as persistent antigenic stimulation together with IL-2 signals results in the upregulation of CD25 and formation of high-affinity heterotrimeric IL-2R. This distinguishes CD8 T cells under persistent antigenic stimulation from naïve and memory CD8 T cells which express heterodimeric intermediate-affinity IL-2Rβγ and rely primarily on homeostatic cytokines (3). Although we cannot directly apply these findings to ongoing human clinical trials due to the differences between mouse surrogate Immuno-STAT vs. CUE-100 molecules in terms of pharmacokinetics, dosage, and treatment regimen, future studies could explore the relative activity and selectivity of reduced Immuno-STAT dose levels and frequencies as it relates to target cell selectivity.

Lastly, modest effects on viral control were observed in these studies, even when combined with PD-1 blockade in chronic LCMV infection. This may be partly due to the nature of the chronic LCMV infection model, representing a stringent challenge with significant systemic viral burden targeting multiple cell types, where targeting one T cell epitope may be insufficient to improve overall viral control despite efficient effector CD8 T cell responses in the targeted T cell population. These considerations are particularly relevant for clinical translation, where diverse strategies are being evaluated, including different IL-2R bias approaches and various targeted strategies directed toward tumors, TME, and specific immune cell populations (3-5).

In conclusion, our study demonstrates the therapeutic potential and feasibility of TCR-directed IL-2 therapeutics for targeting antigen-specific CD8 T cells of interest even in the setting of chronic antigen exposure. Given that numerous clinical trials of IL-2 based drug candidates are ongoing in patients with cancer as well as other indications such as autoimmune diseases, our findings inform both these clinical studies and future preclinical development of IL-2 based agents. Future studies should focus on optimizing IL-2R bias, exploring multi-epitope targeting approaches, and targeted delivery strategies for maximizing therapeutic efficacy while minimizing off-target effects.

## MATERIALS and METHODS

### Mice, virus, and infection

Six- to 8-week-old female C57BL/6J mice were purchased from the Jackson Laboratory (Bar Harbor, ME). Chronically LCMV-infected mice were generated as follows. Mice were transiently depleted of CD4 T cells by injecting them with 300 µg of rat anti-mouse CD4 antibody (GK1.5, BioXCell) i.p. twice on days -2 and 0, followed by intravenous (i.v.) infection with 2 x 10^6^ PFU of LCMV clone 13. For assessing therapeutic effects of Immuno-STAT treatment on LCMV-specific CD8 T cells in the presence of LCMV-specific CD4 T cells, mice were infected with LCMV clone 13 without transient CD4 T cell depletion. Viral titers were determined by plaque assay on Vero E6 cells. All experiments were conducted in accordance with National Institutes of Health and the Emory University Institutional Animal Care and Use Committee (IACUC) guidelines.

### Cell isolation

Spleens were dissociated by passing them through a 70 μm cell strainer (Corning), followed by incubating with ACK lysing buffer (Lonza) and washing with RPMI containing 2% FBS. Livers were perfused with PBS and homogenized via mechanical disruption followed by purification with a 44–67% Percoll gradient (800 g at 20 °C for 20 min). Blood samples were collected in 4% sodium citrate buffer and PBMCs (peripheral blood mononuclear cells) were isolated using lymphocyte separation medium (Corning).

### Reagents, flow cytometry and in vitro stimulations

All antibodies for flow cytometry were purchased from BD Biosciences, Biolegend, or Thermo Fisher Scientific. MHC class I tetramers for H-2D^b^-restricted GP33-41 and GP276-286 epitopes were manufactured from biotinylated monomers prepared in-house, linked to streptavidin-allophycocyanin (APC) or -phycoerythrin (PE) (Thermo Fisher Scientific), for detection of LCMV-specific CD8 T cells. Dead cells were excluded by using the Live/Dead Fixable Blue or Near-IR Dead Cell Stain Kit (Thermo Fisher Scientific). For cell surface staining, antibodies were added to cells at dilutions of 1:20-1:500 in PBS supplemented with 2% FBS and 0.1% sodium azide for 30 min on ice. Cells were washed 3 times and fixed with Fixation/Permeabilization Concentrate of the Foxp3/Transcription Factor Staining Buffer set (Thermo Fisher Scientific), followed by intranuclear staining of transcription factors according to the vendor’s protocols. For detecting cytokine production, 2 × 10^6^ spleen cells were stimulated with GP33-41 or GP276-286 peptide (0.1 μg/ml) in a 96-well round-bottom plate for 5 h at 37 °C in a CO2 incubator in the presence of GolgiPlug (BD Biosciences). Samples were acquired on Canto II (BD Biosciences) or Cytek Aurora (Cytek Biosciences), and data were analyzed using FlowJo v10.10.0 (BD Biosciences).

### Design, Manufacturing and purification of Immuno-STAT proteins

Immuno-STAT protein targeting D^b^GP33-specific CD8 T cells, murine surrogates of CUE-100 molecule with functionally equivalent domains, was generated to assess activity in immunocompetent mice as previously described (11). The murine surrogate retains the design of CUE-100, with an effector attenuated murine IgG2a Fc, an H-2D^b^-GP_33-41_, peptide–MHC complex, and identical human mutant IL-2 components. Human IL-2 is cross-reactive with mouse IL-2R, allowing modeling human IL-2 effects on mouse cells and in mouse models. Immuno-STAT proteins were expressed by stably transfected CHO-K1 cells (ATUM). Proteins were purified from the conditioned media using a two-step method of ProteinA capture with MabSelect SuRe (GE) followed by size exclusion chromatography. For SDS-PAGE analysis, proteins were boiled in SDS sample buffer with or without reducing agent for 5 min before loading 2 µg per gel lane.

### Immuno-STAT therapy, PD-1 blockade, and the combination treatment in vivo

For Immuno-STAT therapy, 15 mg/kg of Immuno-STAT resuspended in PBS with 500 mM NaCl were i.p. injected twice daily for 7-10 days. For PD-1 therapy, 200 μg of rat anti-mouse PD-L1 antibody (10F.9G2, prepared in house) was administered i.p. every 3 days for 7-10 days. Combination treatment was performed by combining these two treatment regimens.

### Analysis of multiparameter spectral flow cytometry

For examining phenotypes of LCMV-specific CD8 T cells, 23-colour spectral flow cytometry data of D^b^GP33- and D^b^GP276-specific CD8 T cells after different treatments were concatenated, and processed for UMAP plugins (nearest neighbors = 15, minimum distance = −0.5 and number of components = 2) (27) and the FlowSOM clustering algorithm (number of meta clusters = 3) (28) using the parameters of TCF-1, SLAMF6, CD73, Ki-67, granzyme B, CD25, CD218a, CD119, T-bet, CD44, Ly-6C, CD69, CX3CR1, CD101, and TIM3 in FlowJo v.10.10.0 (BD Biosciences).

### Statistics

Prism 10.6.1 software (GraphPad) was used for statistical analysis. The differences among the experimental groups were assessed by using a Mann-Whitney test, one-way ANOVA with Dunnett’s correction for multiple comparisons, the Kruskal-Wallis test with Dunn’s correction for multiple comparisons, and two-way ANOVA with Šídák tests for multiple comparisons. The value or the percentage are reported with mean and SD or geometric mean and 95% confidence interval. A p-value of 0.05 or less was considered statistically significant.

## ACKNOWLEDGEMENTS

The authors thank Hong Wu and Chengjing Zhou for their technical assistance, and Wynona Bautista, Nitin Kuman, and Ahmet Vakkasoglu of Cue Biopharma for providing reagents, methods, and input. M.H. holds adjunct positions at Kumamoto University (Japan) and Saitama Medical University (Japan), which are unrelated to the content of this manuscript.

## FUNDING

This work was supported by grants from CUE Biopharma (R.A.).

## AUTHOR CONTRIBUTIONS

M.H. S.N.Q., A.S., and R.A. conceptualized the study and designed the experiments. M.H., M.A.K., Y.Z., N.G., and S.N.Q. performed experiments. M.H., M.A.K., J.N.A. and R.A. analyzed the experiments. G.J.F., Y.Z., N.G., S.N.Q., and A.S. contributed critical materials. M.H. and R.A. wrote the manuscript, with all authors contributing to writing and providing feedback.

## DATA AVAILABILITY

All data are included in the manuscript and supporting information.

## FIGURE LEGENDS

**Supplemental Figure 1. PD-1 and TOX expression in the three clusters of D^b^GP33-specific CD8 T cells during chronic LCMV infection.** Representative histograms showing PD-1 and TOX expression on D^b^GP33-specific CD8 T cells in the three clusters identified in Figure 6. Naïve CD8 T cells (CD44^lo^PD-1^neg^) were included as a control population and are indicated by the grey histogram. Results were pooled from 2-3 experiments with n=1-4 mice per group in each experiment.

**Supplemental Figure 2. Effects of Immuno-STAT alone or in combination with anti-PD-L1 treatment on viral control in a chronic LCMV model with CD4 T cell help.** Chronically LCMV-infected mice (19-25 days post-infection) were left untreated or treated with anti-PD-L1 antibody alone, Immuno-STAT alone, or a combination of anti-PD-L1 antibody and Immuno-STAT for 7-10 days, followed by analysis of viral titers in the spleen and liver. Dotted line indicates the limit of detection. Results were pooled from 5-7 experiments with n=1-5 mice per group in each experiment. Statistical comparisons were performed using the Kruskal-Wallis test with Dunn’s correction for multiple comparisons. Bars and error bars represent the geometric mean and 95% confidence interval. Untx, untreated.

## Notes

Conflict-of-Interest statement: G.J.F has served on advisory boards for iTeos, NextPoint, IgM, GV20, IOME, Bioentre, Santa Ana Bio, Simcere of America, and Geode. G.J.F has equity in Nextpoint, iTeos, IgM, Invaria, GV20, Bioentre, and Geode. R.A. holds patents on PD-1 inhibitory pathway (8,652,465 and 9,457,080) licensed to Roche. Y.Z., N.G., S.N.Q., and A.S. are employed by Cue Biopharma. The remaining authors declare no competing financial interests.

